## Supplementary figures and images for "Maternal High Fiber Diet Protects Offspring Against Type 2 Diabetes"

### Supplementary Figure S1

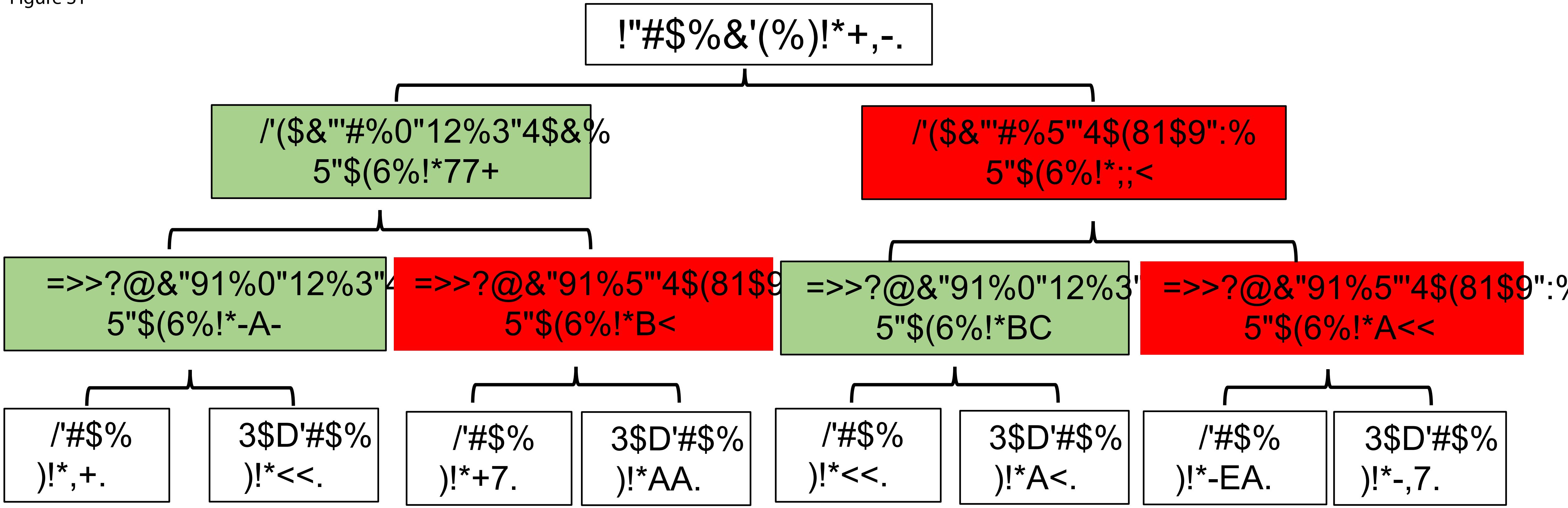

### Supplementary Figure S2

Figure S2

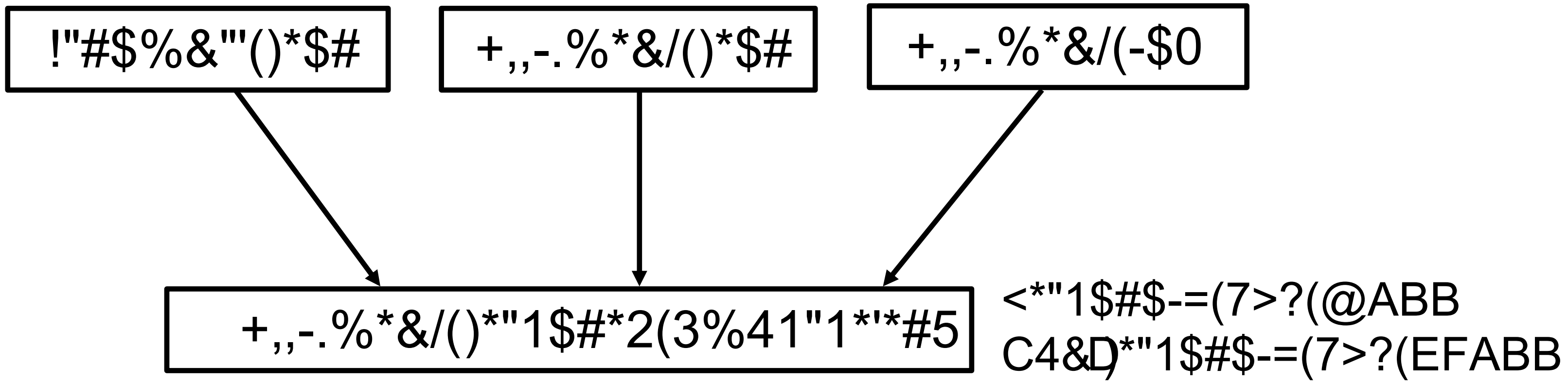


3(67*-8(9(!"#$%&"'()*$#:(+,,-.%*&/()*$#:(+,,-.%*&/(-$0;(

C$#G4%8(-#%H2#H%$=(3%$)$,*&$)
>"5$-*"&(&$#G4%8(."%"I$#$%('$"%&*&/("&)(.%41"1*'*#5(*&,$%$&2$=(7(."28"/$(6J1&'$"%&K;

### Supplementary Figure S3

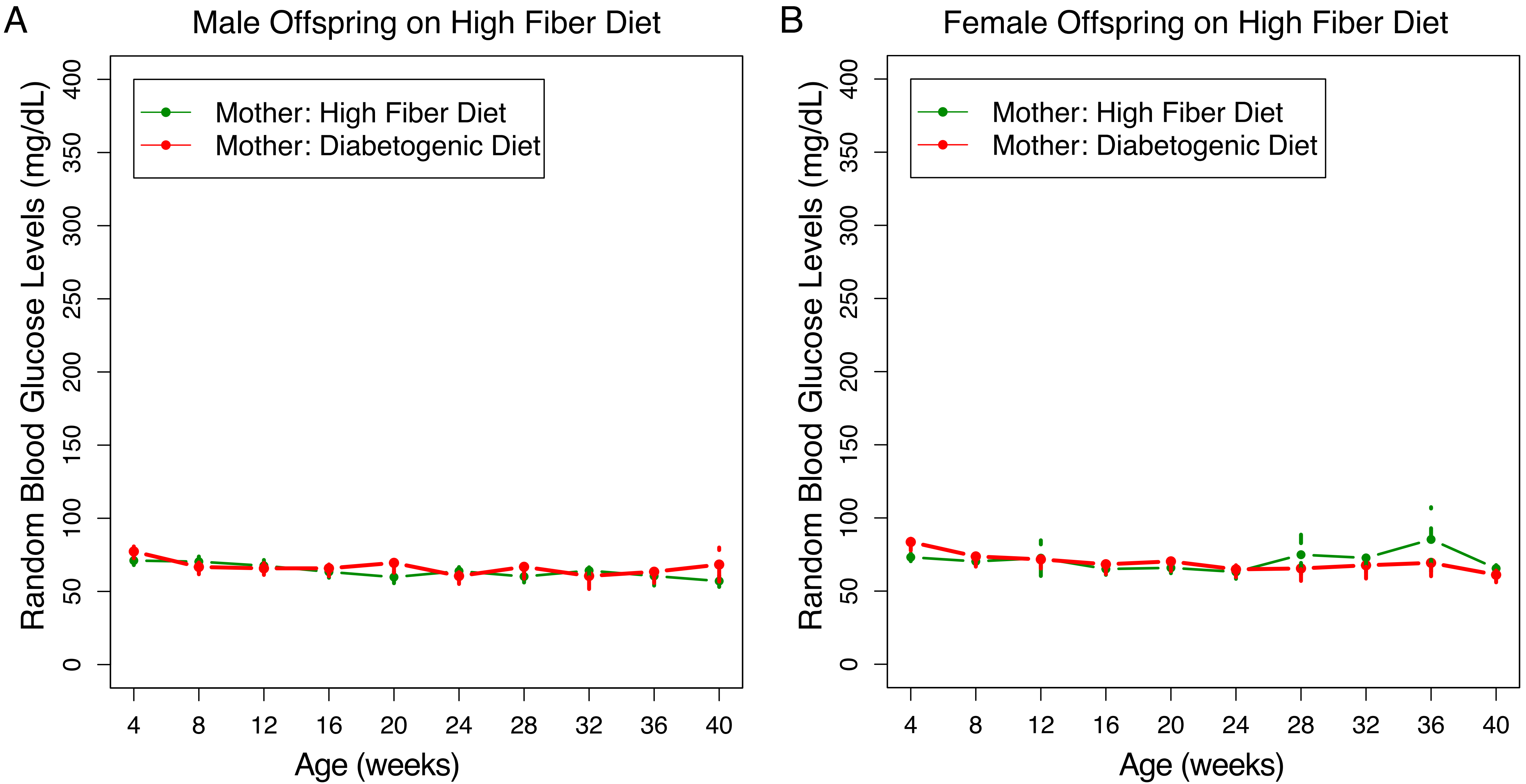
